# Accurate identification of *Helicoverpa armigera–Helicoverpa zea* hybrids using genome admixture analysis: implications for genomic surveillance

**DOI:** 10.1101/2023.11.03.565557

**Authors:** D. Trujillo, T. Mastrangelo, C. Estevez de Jensen, J.C. Verle Rodrigues, R.D. Lawrie, S.E. Massey

**Affiliations:** Department of Agro-Environmental Sciences, University of Puerto Rico - Mayaguez, Mayaguez, PR, USA; Universidade de São Paulo, Centro de Energia Nuclear na Agricultura, 13416-000, Piracicaba, SP, Brazil; USDA-APHIS-PPQ-S&T, 22675 Moorefield Rd, Edinburg, TX 78541, USA; Center for Excellence in Quarantine and Invasive Species (CEQUIS), Estacion Experimental Agricola, Jardin Botanico Sur, San Juan, PR; Department of Biology, University of Puerto Rico - Rio Piedras, San Juan, PR, USA

**Keywords:** *Helicoverpa zea*, *Helicoverpa armigera*, hybrid, F1, admixture, genome, crop pest, genome

## Abstract

*Helicoverpa armigera*, the cotton bollworm moth, is one of the world’s most important crop pests, and is spreading throughout the New World from its original range in the Old World. In Brazil, invasive *H.armigera* has been reported to hybridize with local populations of *Helicoverpa zea*. The correct identification of *H.armigera-H.zea* hybrids is important in understanding the origin, spread and future outlook for New World regions that are affected by outbreaks, given that hybridization can potentially facilitate *H.zea* pesticide resistance and host plant range via introgression of *H.armigera* genes. Here, we present a genome admixture analysis of high quality genome sequences generated from two *H.armigera-H.zea* F1 hybrids generated in two different labs. Our admixture pipeline predicts 48.8 % *H.armigera* for both F1 hybrids, confirming its accuracy. Genome sequences from five *H.zea* and one *H.armigera* that were generated as part of the study show no evidence of hybridization. Interestingly, we show that four *H.zea* genomes generated from a previous study are predicted to possess a proportion of *H.armigera* genetic material. Using unsupervised clustering to identify non-hybridized *H.armigera* and *H.zea* genomes, 8511 ancestry informative markers (AIMs) were identified. Their relative frequencies are consistent with a minor *H.armigera* component in the four genomes, however its origin remains to be established. We show that the size and quality of genomic reference datasets are critical for accurate hybridization prediction. Consequently, we discuss potential pitfalls in genome admixture analysis of *H*.*armigera-H.zea* hybrids, and suggest measures that will improve such analyses.

## Introduction

*Helicoverpa armigera,* the Old World cotton bollworm, is an Old World species of moth, and one of the world’s most important plant pests, whose larvae consume plants belonging to at least 68 plant families (Cunningham and Zalucki 2014). In the New World, *H. armigera* was initially observed in Brazil in 2013 (Czepak et al. 2013), and has subsequently spread throughout much of Latin America (Murúa et al. 2014) (Tembrock et al. 2019), appearing to have undergone multiple introduction events into South America from the Old World (Gonçalves et al. 2019) (Arnemann et al. 2019). There has not yet been a formal identification of *H.armigera* in North America, although it has been intercepted at several ports (Kriticos et al. 2015). The potential economic damage that *H.armigera* could cause in North America is large: $78 billion worth of crops in the United States were estimated to be susceptible to the pest in 2015 (Kriticos et al. 2015).

A closely related species, *Helicoverpa zea*, is native to the New World, and does not have such a wide host range, feeding off over 110 host plants (Kogan et al. 1989). *H.zea* does not possess such a high degree of resistance to common pesticides as that observed in *H.armigera* (da Silva et al. 2020) (although resistance to Bt-proteins has been widely documented in *H.zea* (Burd et al. 2003)), implying it does not pose such an economic threat as *H.armigera*.

*H.armigera* and *H.zea* diverged approximately 1.5 million years ago (Behere et al. 2007), and are able to produce viable hybrids (Laster and Sheng 1995) . *H.armigera-H.zea* hybrids have been reported from Brazil (Anderson et al. 2018) (Valencia-Montoya et al. 2020) (Cordeiro et al. 2020), but have yet to be identified from elsewhere. Adult *H.armigera* are difficult to distinguish from *H.zea* on the basis of morphology, requiring dissection of genitalia (Pogue 2004). Identifying hybrids using such methods is impossible, while larvae of the two species are likewise indistinguishable using morphology (Tay and Gordon 2019). In addition, such methods are inappropriate for screening large numbers of animals. While pure *H.armigera* and *H.zea* can be differentiated using species-specific PCR of the ITS1 region, this method does not work for hybrids (Perera et al. 2015). Hence, genomic methods have great potential utility for accurate species and hybrid identification.

The occurrence of *H.armigera-H.zea* hybrids in Brazil (Anderson et al. 2018) (Valencia-Montoya et al. 2020) (Cordeiro et al. 2020) has implications for pest management programs. Adaptive introgression of genes from invasive pest species into related local species poses a significant threat to global agriculture (Tay and Gordon 2019). A primary reason for studying *H.armigera-H.zea* hybrids in the field is to monitor the adaptive introgression of pesticide resistance genes to *H.zea* from *H.armigera*, which has been subject to intense selective pressure from synthetic pesticides (Walsh et al. 2022). For example, the *CYP337B3* gene, which confers resistance to pyrethroids, has already introgressed into *H.zea* populations in Brazil (Valencia-Montoya et al. 2020). The frequency of pesticide resistance genes in both *H.zea* and *H.armigera* populations has implications for the choice, duration and intensity of pesticide regimens dedicated to their control.

Genes in addition to those responsible for pesticide resistance may also have a propensity to introgress into local *H.zea* populations. For example, *H.zea* lacks genes for gustatory receptors and detoxification compared to *H.armigera*, which may help to explain its more limited range of host plant species (Pearce et al. 2017). These genes may have the potential to introgress from *H.armigera* into *H.zea*, potentially increasing *H.zea’s* agricultural impact by increasing its range of host plants.

*H.armigera* has not yet been formally identified from North America, partly due to difficulties in distinguishing the species from *H.zea*. *H.armigera* was reported in Puerto Rico in 2014 and 2018, however since that time has not been reported again (Flores-Rivera et al. 2022) The Caribbean represents a major transit route for pests and pathogens between North and South America (Waugh 2009), forming a ‘Caribbean corridor’, so Puerto Rico is a critical location for monitoring the potential spread of *H.armigera* from the South American continent into North America.

In this study, we implemented a bioinformatic pipeline to predict hybridization proportions by using whole genome sequences. We used the genomes of two lab generated *H.armigera-H.zea* F1 hybrids to confirm the accuracy of our admixture analysis procedure. We demonstrate that genomes from Puerto Rican and North American *H.zea* genomes generated as part of the study do not show evidence of hybridization with *H.armigera*. However, four attributed North American *H.zea* genomes from a previous study displayed potential evidence of hybridization, representing the potential early presence of *H.armigera* in North America. We show that high quality genome sequence data, reference genomic datasets and careful SNP filtration approaches are important for the accurate determination of hybridization proportions.

## Methods

### Collection and maintenance of parental species

Individual *H.zea* animals were collected by USDA APHIS collaborators (Todd Gilligan) and shipped to our lab in San Juan in ethanol from Colorado in 2015 (HzCol), Illinois in 2016 (HzIll), Maine in 2016 (HzMaine) and North Carolina in 2016 (HzNC). Species identifications were performed using species specific PCR of the ITS1 region, following the methods of Perera et al., 2015.

All live *Helicoverpa* colonies were maintained under the following conditions: 25 ± 2 °C, 57±9 % relative humidity, photoperiod of 15 hours of light and 9 hours of dark (15: 9 LD). Female pupae were placed in incubators at 22.7 ± 1.6 °C, 82 ± 4 % relative humidity, photoperiod of 15:9 LD, females were placed at a lower temperature to synchronize the emergence of adults with males (Armes et al, 1992; Colvin & Cooter, 1994). The larvae were fed with Gypsy Moth Diet (Frontier Agricultural Sciences, Product # F9630B, Newark DE): 140.2 g of dry mix, 20 g of fats and sugars, 1.6 g of vitamin mix, 0.8 g of aureomycin, 1000 ml of distilled water, with the addition of 12 ml of formaldehyde 1%, and 2.5 g of FABCO mold inhibitor (Frontier Agricultural Sciences, Product # F0018, Newark DE); the agar was dissolved, when the temperature was ∼50°C the rest of the reagents were added. Each larva was maintained in transparent plastic cups of 30 ml containing diet. The pupae were maintained in the same cups.

Emerged adults and pupae near to emergence were placed in white plastic buckets of 18.9 l, the upper part of the buckets was covered with cheesecloth (DeRoyal, BIDF2012380-BX, Tennessee) for oviposition. Inside each bucket a Petri dish with autoclaved sand a potted tomato plant was placed to increase relative humidity. The adults received the following diet recipe modified from Grzywacz et al. (2002): 500 ml of distilled water, 50 ml of honey, 10 ml of solution 28 % of Vanderzant vitamin mixture (Sigma, V1007, USA), 1 g of methyl-4-hydroxybenzoate (Sigma, H3647, USA), and 1 ml of ethanol 95 %; methyl-4-hydroxybenzoate was dissolved in the 95% ethanol, then all the ingredients were mixed in the water and fed to adult moths using cotton wicks. The cheesecloth with the oviposited eggs was placed in Ziploc bags of 3.8 liters with fine strips of larval diet. Once larvae emerged, they were transferred to cups with diet. Prior to molecular work, all samples were stored in 90% ethanol in a –20 °C freezer until DNA extractions were performed.

In CEQUIS, separate colonies of *H.armigera* and *H.zea* were maintained. The colony of *H.armigera* was obtained from five larvae and 30 pupae from Brazil courtesy of Dr Thiago Mastrangelo, University of Sao Paolo. The insects were collected from Bahia (12°13’53’’S, 45°44’44’’W) in 2016 and were introduced to quarantine facilities of the Center of Excellence in Quarantine & Invasive Species on February 4, 2017, under Puerto Rico Department of Agriculture Permit number OV-1617-03 and USDA-APHIS Permit number P526P-15-04600 to Dr. José Carlos Verle Rodrigues. The initial colony of *H.zea* was obtained from larvae collected in Isabela, Puerto Rico, from pigeon peas on November 11, 2015. During the F9 generation, a reintroduction of insects was done, from larvae collected in Isabela in corn on November 22, 2016.

### Breeding of the hybrids

The first hybrid included in the study (PRh) was generated in our lab from a male *H.armigera* from Brazil, and a female *H.zea* from Puerto Rico. The resulting hybrid animal was a female. Using the same rearing methods described above, 15 *H.zea* female pupa and 15 *H.armigera* male pupa were placed into a white plastic bucket with cheesecloth lid and allowed to emerge, mate, and oviposit. All surviving F1 hybrids resulting from this cross were labeled and stored in a –20°C freezer.

Genome sequences were generated from parental animals. A sequence from a male *H. armigera* from Brazil (HaM) was generated. This animal was an adult male *H.armigera* from the *H.armigera* colony initiated in CEQUIS, and was one of the parents for the F1 hybrids generated in the lab. A sequence from a female *H.zea* from Puerto Rico (HzF) was also generated. HzF was reared following the conditions described above, and was a parent for the *H.armigera*-*H.zea* F1 hybrids (PRh in this study) generated in the lab.

The second hybrid included in the study (MAh) was generated from a female *H.armigera* from Spain and a male *H.zea* from the mainland USA by the USDA APHIS Otis Lab in Buzzards Bay, Massachusetts in 2017 by Dr. Hannah Nadel. The Spanish *H.armigera* mother was from a colony maintained in Spain, however the original collection was from Portugal. The *H.zea* father used in the MAh cross were supplied by Benzon Research Inc. (Carlisle, PA, USA). This hybrid was reared under the same rearing conditions described previously. Both MaH and PRh hybrids were females.

### DNA extraction and sequencing

DNA samples were obtained from the animals using QIAGEN blood and tissue DNA extraction kits (QIAGEN INC., Cat No,/ID 69506) following the manufacturer’s protocol, with the exception of the Colorado, Illinois and North Carolina samples, which were extracted using the CTAB method (Calderón-Cortés et al. 2010). DNA quality was assessed using a NanoDrop 2000 (ThermoFisher Scientific, Waltham, MA) to assess DNA concentration (ng/uL) and absorbance (A260/280) and gel electrophoresis (1.5 % agarose) to assess integrity and molecular weight. After checking DNA concentration and quality, the eight samples were shipped overnight on ice to the Rapid Genomics sequencing laboratory in Florida (www.rapid-genomics.com).

Paired end sequencing was conducted by RAPID Genomics on the Illumina HiSeq-X platform (sequencing statistics are displayed in Supplementary Table 1). The sequence data has been deposited in the National Center for Biotechnology Information (NCBI) Short Read Archive (SRA) under the Accession numbers SAMN35038651 (PRh), SAMN35038652 (MAh), SAMN35038653 (HzF), SAMN35038654 (HaM), SAMN35038647 (HzCol), SAMN35038648 (HzIll), SAMN35038649 (HzMaine), SAMN35038650 (HzNC). Additional genomic data was used in the analysis, consisting of 29 *H.armigera,* 9 *H.zea* and 9 *H.armigera*-*H.zea* hybrids, from (Anderson et al. 2018) (Table 1). Raw sequence data for these animals were obtained from the Commonwealth Scientific and Industrial Research Organisation (CSIRO; https://data.csiro.au/collection/csiro:29053). Genome sequence analysis was performed on an Amazon Web Services c6g.4xlarge instance (comprising the AWS Graviton2 processor, 16 vCPUs, 32 Gb memory and Amazon Linux platform).

**Table 1.**
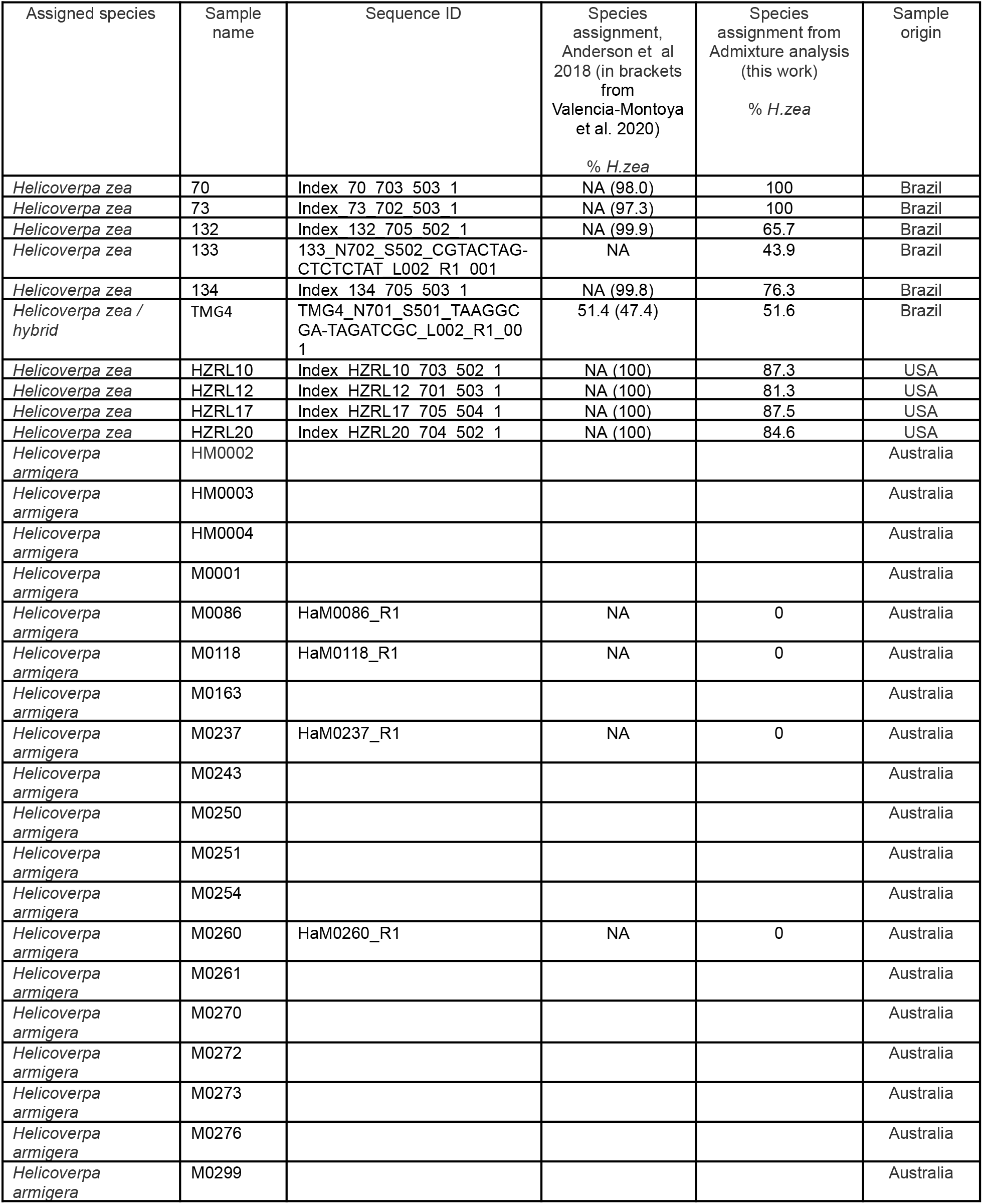
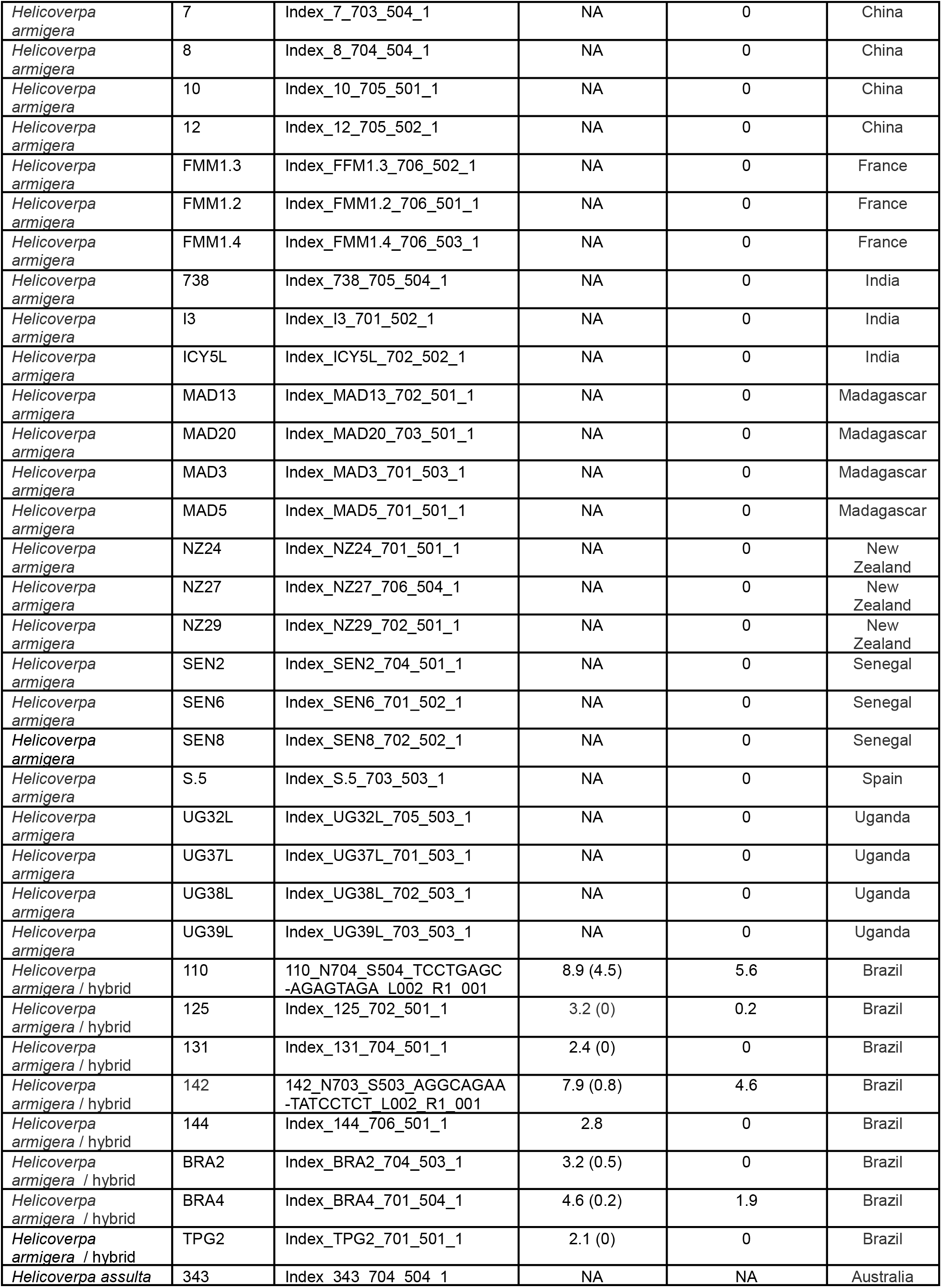
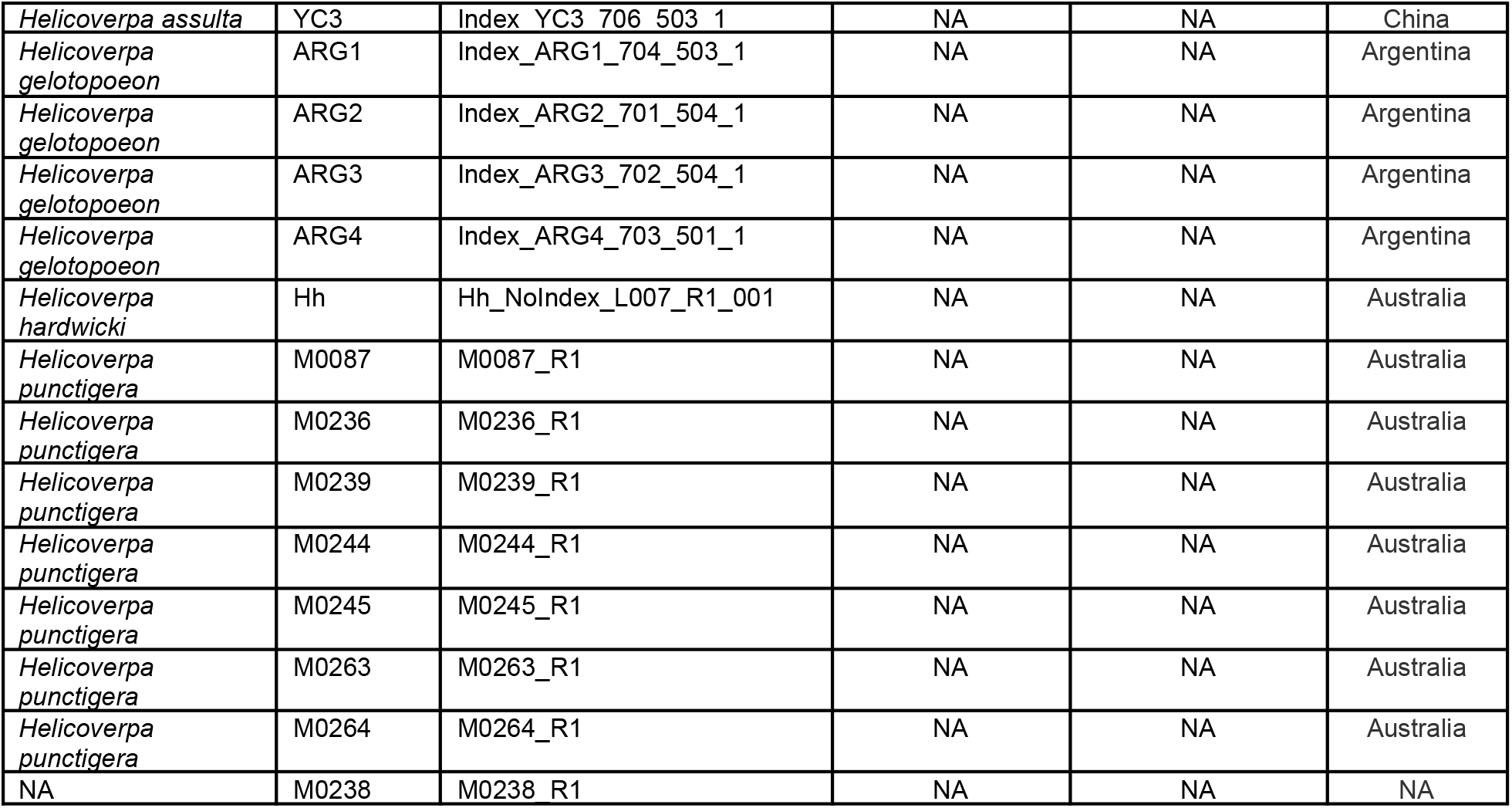
Details of 47 additional *Helicoverpa* genomes used in the Admixture analysis. Genome data was obtained from (Anderson et al. 2018). ‘NA’ means ‘not available’.

### Mapping and SNP calling procedure

Using fastp (Chen et al. 2018), sequences were removed if they did not fulfill the criteria of 95% nucleotides > Q20, 3’ trimming was conducted by quality, and polynucleotide runs (6 or more consecutive). Filtered and trimmed sequences were repaired using the repair.sh script of BBMap (v37.99) (sourceforge.net/projects/bbmap). They were then mapped to the *H*.*armigera* reference genome (Pearce et al. 2017) (all-chr-r.fasta, obtained from CSIRO at https://data.csiro.au/collection/csiro:29053v1), using BBMap in paired-end mode. The resulting sam files were converted to bam files, and sorted using SAMtools (Danecek et al. 2021).

BCFtools mpileup (Danecek et al. 2021) was used for variant calling. Bam files for all *Helicoverpa* genomes in the study were processed together, to improve the accuracy of calls of SNPs shared across genomes. After SNP calling, the resulting vcf files were filtered using vcftools (Danecek et al. 2011), removing those SNPs that possessed mean read depth (min-meanDP < 5), Q value (Q < 20) and minor allele frequency (MAF < 0.05).

### Admixture analysis

Admixture v1.3.0 (Danecek et al. 2021) was used for genome admixture analysis. Sex chromosomes (chromosome 1) were excluded from the analysis. Plink (Purcell et al. 2007) was used to convert the combined vcf file into bed format, which was used as input for the Admixture analysis, which was run using K=2. Admixture output was visualized using the R ggplot2 package.

### Identification of Ancestry Informative Markers

34 *H.armigera* and 7 *H.zea* genomes were identified using the unsupervised clustering approach of Admixture, described above. The genotype data from these genomes was then used to identify SNPs that possessed a minor allele count (MAC) of 7 for the *H.zea* genomes and 1 for *H.armigera* genomes, using vcftools. These were then pruned by removing all SNP positions, where a SNP was completely absent (GT = 0/0) from one or more *H.zea* genome.

## Results and Discussion

Genome sequencing results are shown in Table 2, and show that the quality of the raw sequences was high for all eight genomes. For consistency, SNP calling was jointly conducted on the *Helicoverpa* raw sequence reads generated by (Anderson et al. 2018), and on the sequences generated as part of this study. Filtering resulted in the removal of a large proportion of SNPs (83 %); this might be reduced in future by increasing sequence coverage in the overall dataset.

**Table 2.** Mapping and SNP statistics, and hybridization proportions, of eight new genome sequences generated in the study.

| Sample name | Average genome sequencing depth | Number of SNPs after filtering | Species assignment from Admixture analysis (this work) % *H.zea* | Geographic origin |
| --- | --- | --- | --- | --- |
| PRh | 31.4 | 9230976 | 51.2 | F1 hybrid of male *H.armigera* from Brazil and a female *H.zea* from Puerto Rico |
| MAh | 34.7 | 8990547 | 51.2 | F1 hybrid of a female *H.armigera* from Spain and a male *H.zea* from USA |
| HaM | 43.7 | 7189237 | 0 | Brazil |
| HzF | 37.4 | 5139536 | 100 | Puerto Rico |
| HzCol | 25.0 | 5185015 | 100 | Colorado, USA |
| HzIll | 25.0 | 5190809 | 100 | Illinois, USA |
| HzMaine | 25.3 | 5188817 | 100 | Maine, USA |
| HzNC | 26.0 | 5793330 | 100 | North Carolina, USA |

### Predicted hybridization proportion of the two lab-reared F1 hybrids

In the hybrid animals, approximately equal proportions of the genome originate from both *Helicoverpa* species (51.2 % *H.zea* : 48.8 % *H.armigera* in both PRh and MAh). These data are displayed on the Admixture plot (Figure 2). In both cases, the Admixture prediction is not exactly 50% *H.zea* : 50% *H.armigera* for either hybrid, even though in the case of PRh, genomes derived from the parental populations were 100% *H.zea* (HzF) and 100% *H.armigera* (HaM). This may be due to two reasons. Firstly, the (male) parental insect population may have possessed a degree of hybridization because they were collected originally from Brazil near where early hybrids have since been detected (Valencia-Montoya et al., 2020). However, it is notable that the MAh F1 hybrid also has the same *H.armigera* : *H.zea* ratio (48.8 % : 51.2 %). This would mean that the *H.zea* from the USA, used to generate the hybrid would also have to have had a low level of *H.armigera* admixture; this seems more unlikely than for an *H.zea* insect from Brazil, where the presence of hybrids has been validated.

**Figure 1.**
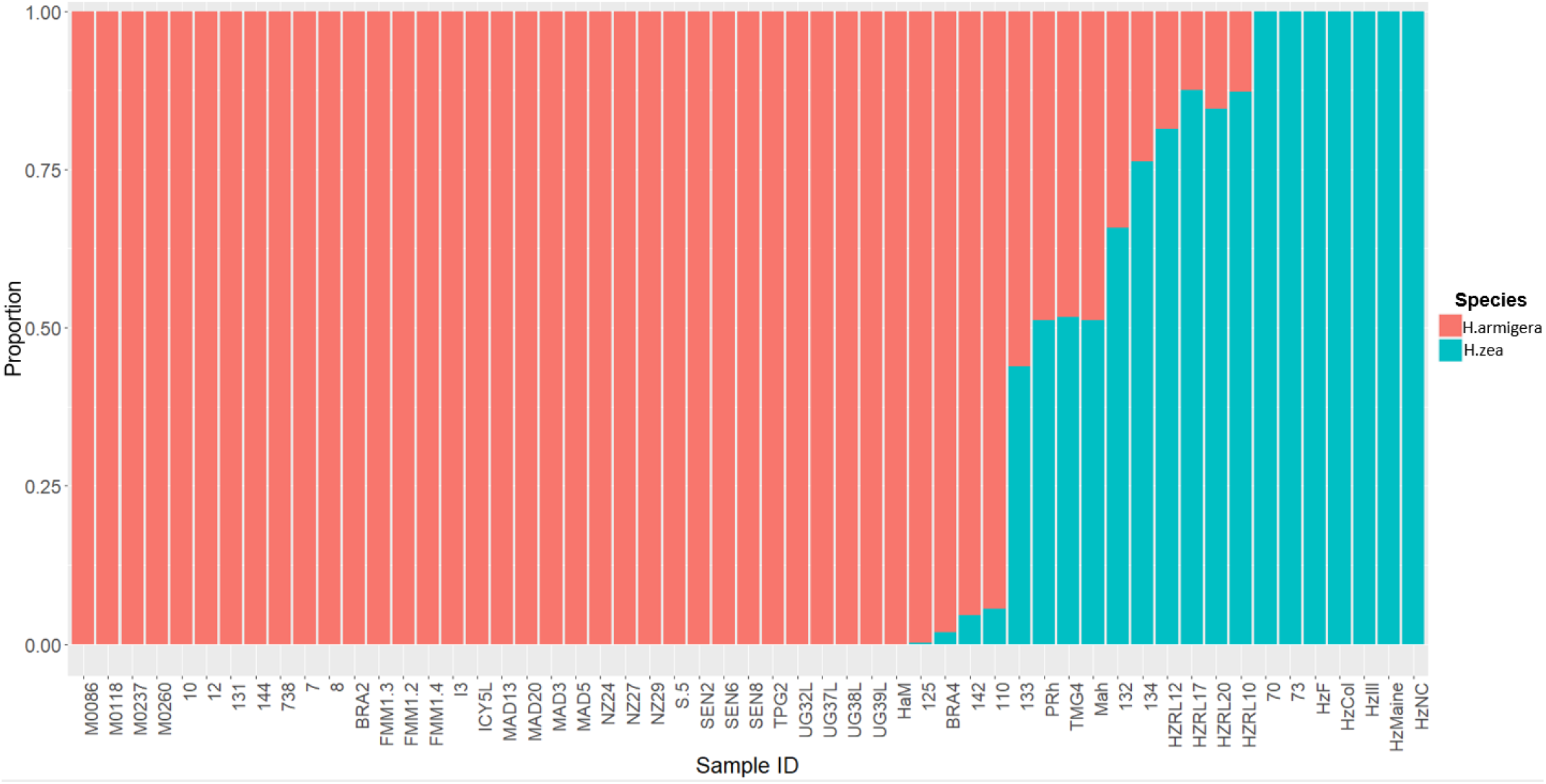
Admixture analysis of *Helicoverpa armigera* and *Helicoverpa zea* genomes The bar plot shows the relative proportions of *H.armigera* and *H.zea* present in *Helicoverpa* genomes generated in this study (PRh, MAh, HaM, HzF, HzCol, HzIll, HzMaine, HzNC), and from (Anderson et al. 2018). The Admixture analysis used K=2, and excluded sex chromosomes.

Secondly, the Admixture analysis may lack exact precision. This may be the result of a limited number of pure *H.zea* in the dataset (seven), which means that the genetic diversity of the species is not adequately represented.This is supported by the observation that the *H.armigera* : *H.zea* ratio is the same for both F1 hybrids: this indicates a systemic bias in admixture prediction.

Encouragingly, even though *H.armigera* and *H.zea* are closely related species, the Admixture analysis is capable of accurately identifying the relative proportions present in an F1 hybrid genome. In future, accuracy may be improved by refinements in SNP calling, increasing the sequencing depth in the overall dataset and adding additional genomes, particularly from *H.zea*. Admixture analysis may be affected by a small sample size of one or more of the reference populations (Lawson, van Dorp, and Falush 2018). In the analysis, even after the addition of five *H.zea* genomes generated in this study, only seven non-hybridized *H.zea* genomes were apparent.

### Predicted hybridization proportions of other Helicoverpa spp. genomes

From the new genome data generated by the study, the analysis indicated that the animals identified as *H.armigera* (HaM), and *H.zea* (HzF, HzCol, HzIll, HzMaine, HzNC), were non-hybridized animals. All Old World *H.armigera* datasets from (Anderson et al. 2018) were identified as non-hybridized, as expected.

The Admixture analysis reveals some discrepancies with those previously published for 47 previously sequenced *Helicoverpa* genomes (Anderson et al. 2018), which were used as a reference dataset here and in other studies. Most of the animals previously identified as 100 % *H.zea* (Anderson et al. 2018; Valencia-Montoya et al. 2020) are predicted in our analysis to have a *H.armigera* component (132, 133, 134, HZRL10, HZRL12, HZRL17, HZRL20), while several specimens previously identified as hybrids (Anderson et al. 2018; Valencia-Montoya et al. 2020) were identified here as 100% *H.armigera* (131, 144, BRA2, TPG2) (Table 1). TMG4, previously described as a *H.zea* hybrid (Anderson et al. 2018; Valencia-Montoya et al. 2020), is also predicted by our analysis as a hybrid and appears to be F1, given its predicted proportion of 48.8% *H.armigera*. Given that this animal was collected in August 2013, this implies that hybridization occurred one generation previous to the collection date.

A key difference between our study and (Anderson et al. 2018; Valencia-Montoya et al. 2020) is that our inclusion of two lab-reared F1 hybrids allows us to verify the accuracy of our analysis. Potential explanations for differences in predicted hybridization proportions reported in (Anderson et al. 2018) may include lack of filtration after SNP calling, and the lower number of *H.zea* in the dataset (leading to a limited reference population for this species). In addition, in (Anderson et al. 2018) SNPs were called on a dataset which included *Helicoverpa punctigera*, *Helicoverpa gelotopoeon*, *Helicoverpa hardwicki* and *Helicoverpa assulta*. In our method, our simultaneous SNP calling procedure only included *H.armigera* and *H.zea* datasets. In addition, in our analysis we chose not to include the Z sex chromosome (chromosome 1), focussing only on autosomes.

The reason for differences between our study and the predicted species proportions described in (Valencia-Montoya et al. 2020) (Table S4, Valencia-Montoya et al; Table 1 this study) is less clear, given that the authors used filtration criteria similar to our own, and only called SNPs against *H.armigera* and *H.zea* genomes, rather than including additional *Helicoverpa* spp. in their analyses. However, the ancestry proportions that they report in Table S4 are derived from ∼1 million SNPs identified as segregating between the two species, whereas we base our ancestry proportions on Admixture analysis, consequently methodological differences may provide the source of the discrepancy.

### Potential H.armigera hybridization detected in North American H.zea from 2005

The identification in the reference dataset of potential *H.armigera*-*H.zea* hybrids from North America (HZRL10, HZRL12, HZRL17, HZRL20), with predicted *H.armigera* proportions of 12.7 %, 18.7 %, 12.5 % and 15.4 %, respectively (Table 1) is interesting, given that *H.armigera* has not been formally identified in the mainland US, and that *H.armigera* was first detected in the Americas in 2013 in Brazil (Czepak et al. 2013). This may therefore represent an early presence of *H.armigera* in the Americas.

The samples were originally described in a 2007 study that constructed a phylogeny of *Helicoverpa* spp. using mitochondrial DNA (Behere et al. 2007), and their genome sequences, used in the study described here, were described in (Anderson et al. 2018). The samples are recorded as having been collected from ‘Riverland, NY’ (Anderson et al. 2018), however this location is unclear. Dr Daniel Gilrein supplied the *H.zea* samples (Behere et al. 2007), and is based at the Long Island Horticultural Research and Extension Center (LIHREC), Riverhead, NY. The origin of the samples is confirmed as Riverhead, NY (personal communication, Dr Dan Gilrein).

The four samples were collected in 2005, in September / October (personal communication, Dr Dan Gilrein). Significantly, this date predates the first reports of *H.armigera* in the New World in 2013 in Brazil (Czepak et al. 2013). In order to confirm this result, 8511 AIMs were identified, as described in Methods. Unsupervised clustering allowed the *a priori* identification of 34 *H.armigera* and 7 *H.zea* non-hybridized genomes (Table 1). These were used to identify SNPs that preferentially segregate in one species or the other (AIMS). The 8511 AIMS thus identified indicate a *H.armigera* component ranging from 25.8 to 31.1 % in the four genomes (Table 3). The predicted presence of a *H.armigera* component is consistent with the results from the Admixture analysis.

**Table 3.**
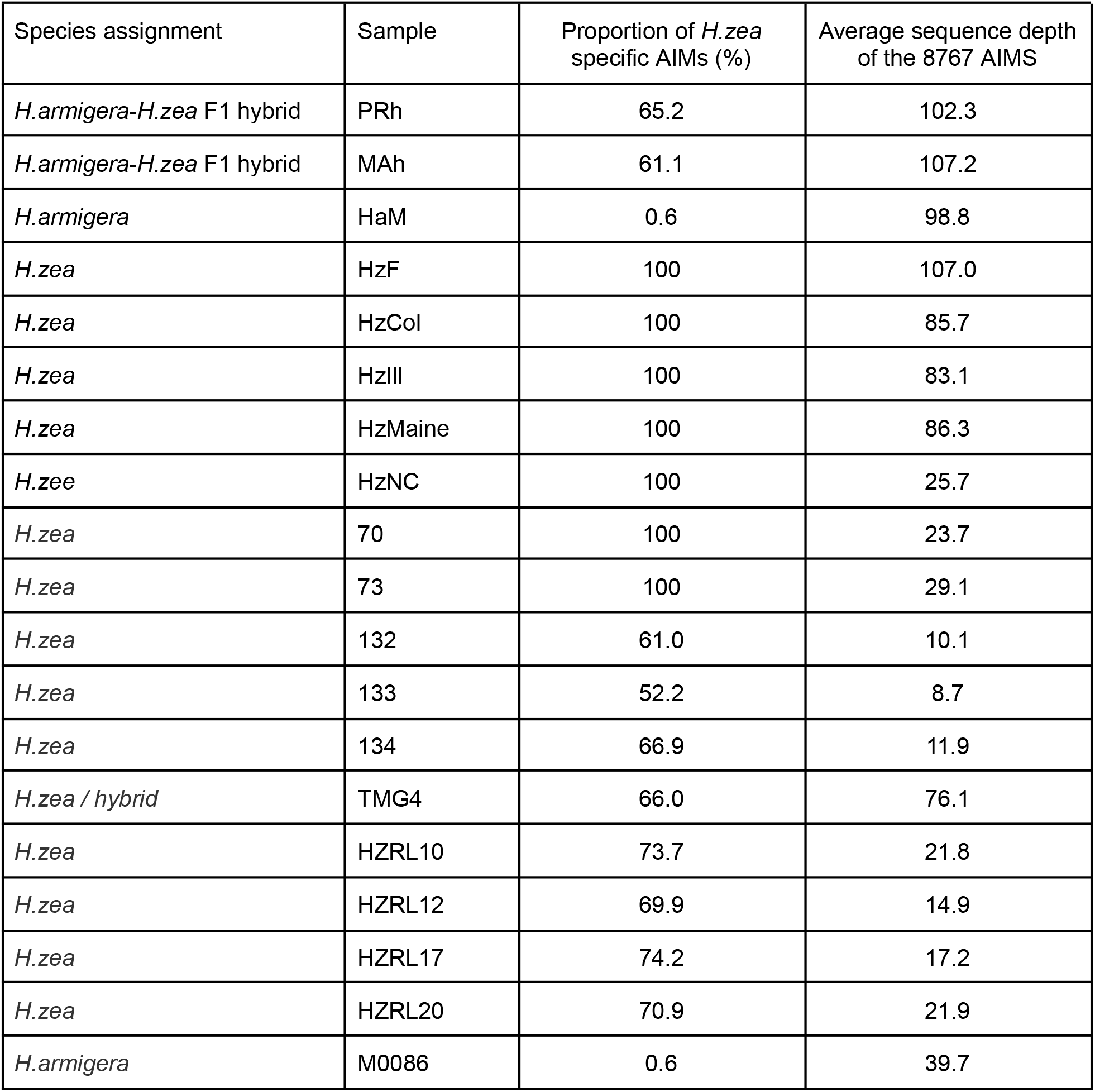
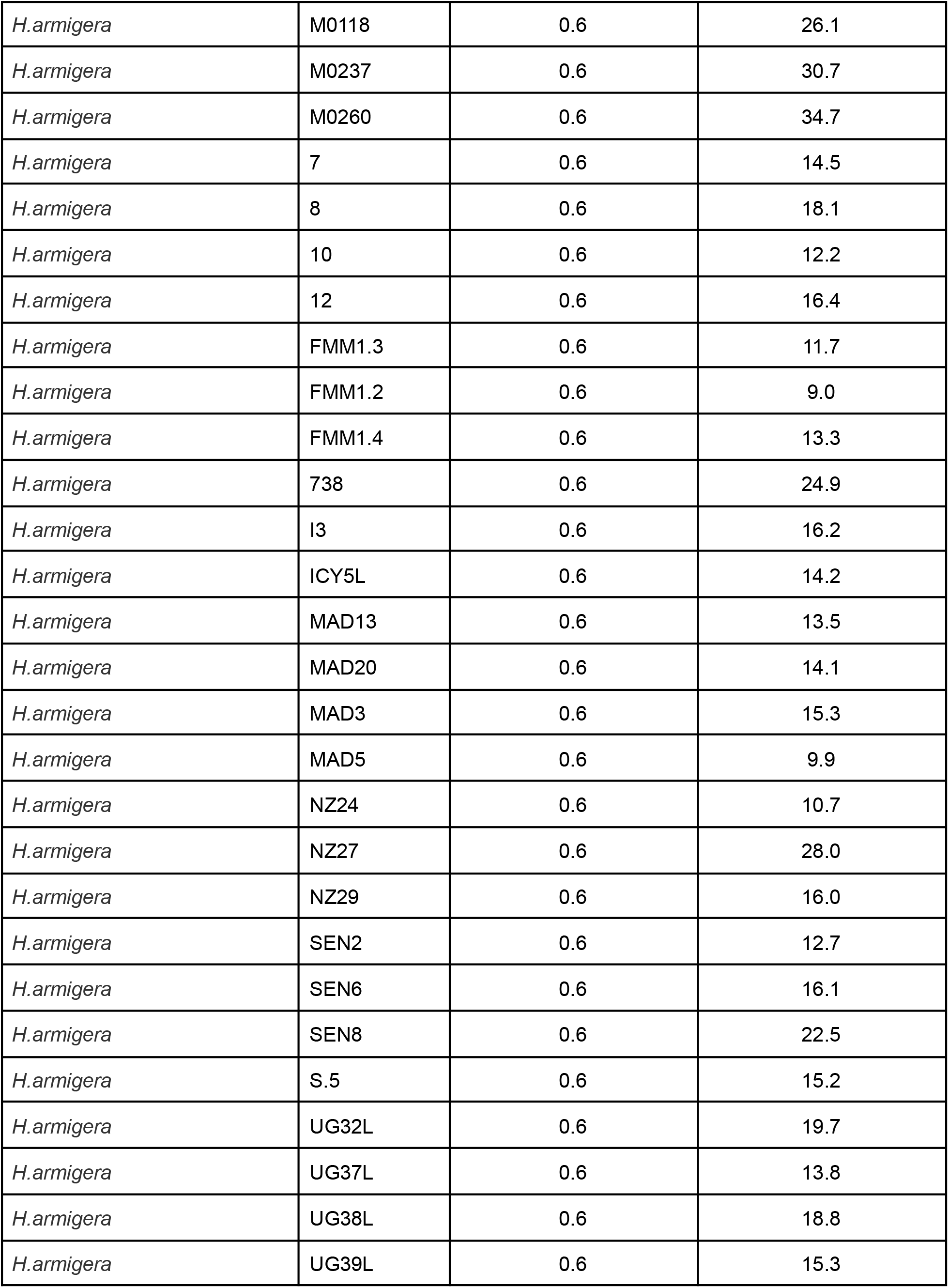
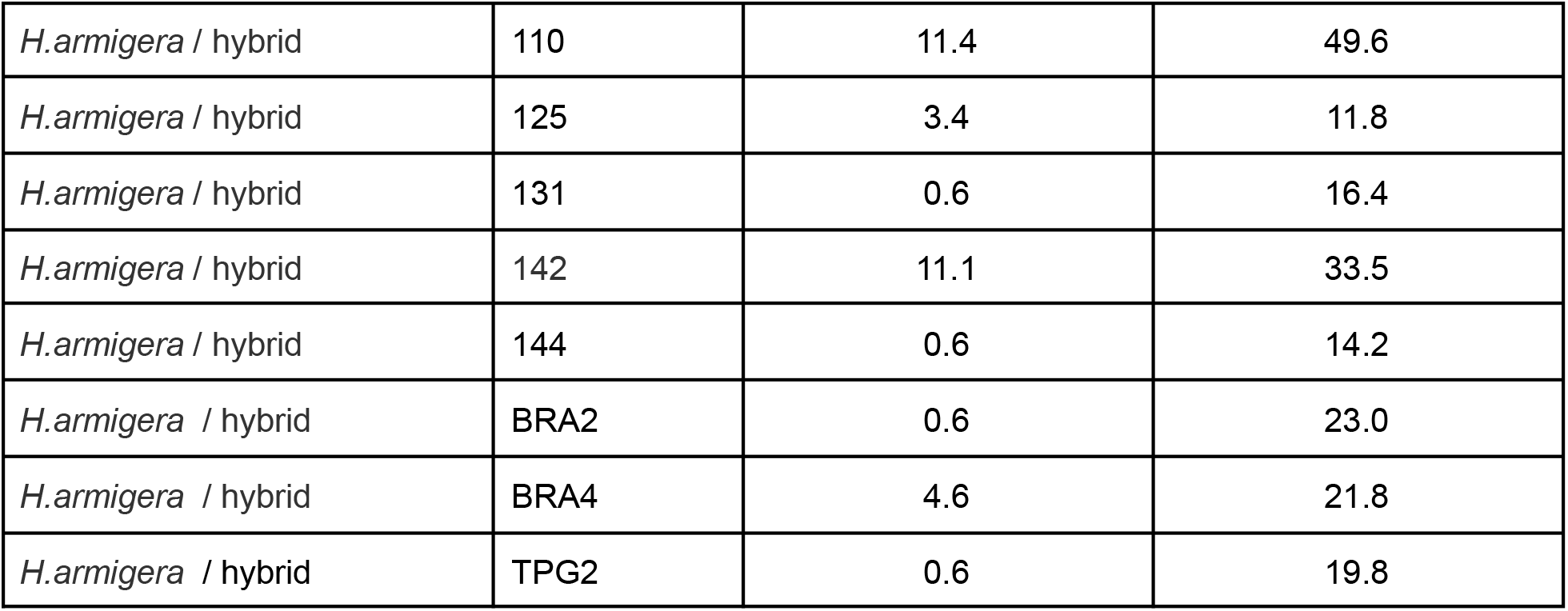
Proportion of *H.zea* AIMs present in different genomic datasets 8511 AIMs were identified as described in Methods. The proportion of *H.armigera* - specific AIMs identified in the different genomes is listed. The proportions were determined by comparison with the filtered SNPs produced from the SNP calling procedure described in Methods.

Regarding the accuracy of this approach, using a reduced set of SNPs is not expected to give the same accuracy as the whole genome considerations utilized by Admixture, however the unsupervised clustering approach represents an independent manner of assessing a potential *H.armigera* contribution to *H.zea* genomic datasets. The *H.armigera* component is higher than predicted by the Admixture approach, which gives 12.5 to 18.7 % *H.armigera*. Notably, the predicted *H.armigera* proportion for the two F1 hybrids is 34.8% (PRh) and 38.9% (MAh) (Table 3), which underestimates the true proportion of 50%. One potential source of error is uneven distribution of AIMs along the chromosomes. Another is that the *H.zea* dataset was limited in size, and so this reduces the accuracy in identifying species-specific AIMs. The low level (0.6 %) of *H.zea* AIMs detected in most of the *H.armigera* genomes reflects the AIM selection approach: the *H.zea* AIMs were present in all 7 *H.zea* genomes, and were also found to be present in at most one *H.armigera* genome in the reference dataset.

Finally, it is possible that low sequencing depth may affect the predicted hybridization proportions. Given that the SNPs are called against a *H.armigera* reference genome, then if a SNP position has low or no read depth in a particular genome, the SNP calling software will call the *H.armigera* genotype at that position. This means a bias toward calling *H.armigera* AIMs when sequencing depth is low. For example, HZRL10 has 333 AIM positions where there is no sequence coverage, reflecting its low average sequencing depth of 21.8 for the AIM positions. In total there are 986 AIM positions where DP < 5, and so cannot be called with confidence; these constitute 11 % of the total number of AIMs. The AIM positions where there is no sequence coverage are by default identified as *H.armigera* (reflecting the reference genome sequence at those positions). This therefore can account for a proportion of *H.armigera* AIMs in the HZRL10 genome sequence, but not all. This observation may also account for a proportion of the *H.armigera* ancestry in HZRL10 detected by the Admixture analysis.

Further work will be required to validate or discount these observations. In particular, the approaches described are not able to distinguish sample contamination from hybridization. Larger, high quality datasets will be necessary in order to distinguish these two alternative scenarios. Development of such fine-grained methods will have value in screening of historic samples and detection of contamination in hybridization studies, which is currently difficult to detect (a method developed by SEM for detecting contamination of NGS datasets, mitoscan https://github.com/semassey/Scanning-NGS-datasets-for-mitochondrial-and-coronavirus-contaminants/blob/main/mitoscan.sh, maps reads against all NCBI mitochondrial genomes, however it is not able to distinguish contamination by closely related species, due to cross-mapping between closely related mitochondria).

### The use of genome admixture analysis for the identification and control of Helicoverpa infestations

We have shown the efficacy of genome admixture analysis for verifying the identity of *Helicoverpa* hybrids, which are morphologically cryptic, and so recalcitrant to traditional identification methods, as is the identification of the two *Helicoverpa* species themselves. We found that increasing the number of *H.zea* genomes in the analysis improved the accuracy of admixture prediction, for the *H.zea* and *H.armigera* genomes, and the two F1 hybrid genomes generated in the study. Likewise, filtering based on sequencing depth also had a similar effect, although we were restricted in increasing filtering stringency, given limitations in sequencing depth in the dataset. Future improvements in accuracy will arise from greater average sequencing depth in the reference genomes used in admixture analyses. Finally, for accurate hybrid identification, whole genome approaches are most likely to yield the precision necessary for understanding the dynamics of *H.armigera* invasivity in the field.

In addition to the indirect detection of *H.armigera* in a region via identification of *H.armigera*-*H.zea* hybrids, determining the presence of the hybrids will have utility for monitoring the occurrence and spread of pesticide resistance. This is desirable because *H.armigera* populations in the Old World have typically been subjected to significant pesticide exposure, thus leading to the evolution of resistance (Valencia-Montoya et al. 2020). Hybridization with local *H.zea* populations is expected to lead to the introgression of pesticide resistance genes from the *H.armigera* genomic component (Valencia-Montoya et al. 2020). The phenomenon of rapid introgression of pesticide resistance genes between sister species has been observed in *Anopheles* spp. exposed to selection pressure from pesticide exposure (Norris et al. 2015). The evolutionary dynamics would be expected to be rather similar in crop pests such as *Helicoverpa* spp.

Host plant preference is another agriculturally relevant phenotype that may be influenced by hybridization and gene introgression is that of host plant preference. *H.armigera* has a considerably more extensive plant host range than *H.zea*, apparently partly due to its larger number of gustatory receptor and detoxification genes compared to *H.zea* (Pearce et al. 2017). Adaptive introgression of these genes from *H.armigera* into local populations of *H.zea* may cause changes in the host plant preferences of *H.zea*, a process consistent with the ‘hybrid bridge’ hypothesis of host shifting of herbivorous insect pests (Floate and Whitham 1993). Furthermore, increasing ease of *H.armigera-H.zea* hybrid detection will allow for the collection of empirical evidence for whether hybridization will influence changes in pesticide susceptibility or feeding behavior. Currently, because hybrids are extremely difficult to identify, empirical data for these phenotypic changes are near impossible to collect.

Puerto Rico is a stepping stone between North and South America, given its geographic location and possession of a major port in San Juan, through which agricultural produce enters and exits the United States. This transit route for agricultural pests and pathogens comprises part of a ‘Caribbean corridor’. So far, there are no reports in the literature on sustained *H.armigera* populations in North America or Puerto Rico. One potential route for the spread of *H.armigera* into North America from South America may be through Puerto Rico.

The detection of *H.armigera-zea* hybrids can reveal aspects of the population dynamics of both species and help inform control strategies. The accurate determination of hybrid proportions can also indicate whether species boundaries are maintained, given that hybridization is often maladaptive.

Accurate admixture prediction methods for *Helicoverpa* species are essential for the design of accurate high throughput hybrid identification tools, and so the datasets generated as part of this study will be useful in the development of tools for the rapid, economical and accurate identification of pure species or hybrids. Future detection of hybrids from Puerto Rico and potentially North America will help inform control regimens, facilitated by the development of rapid molecular tests to accurately determine hybrids. In particular, if there is detection of *H.armigera* in North America, screening of local *H.zea* populations for hybridization could be used to assess whether breeding has occurred.

## Acknowledgements

This work was supported by USDA/APHIS Agreement AP20PPQS&T00C161. It may not necessarily express APHIS’ views. We would like to thank Dr Todd Gilligan, USDA-APHIS, for providing USA specimens, Dr Hannah Nadel, Otis Laboratory MA, for providing the MAh hybrid, Dr Fernando Rodrigues da Silva, University of Florida, for support during rearing insect colonies, and Patricia Caligari, UPR-CEQUIS, for assistance during DNA preparations for sequencing. We would also like to thank Dr Tom Walsh and Dr Tek Tay (CSIRO), and Dr Dan Gilrein (LIHREC, Cornell University, Riverhead, NY) for valuable discussion regarding the specimens collected from NY state. Lastly, we thank an anonymous researcher for helpful comments prior to submission.

## Authors contributions

Conceptualization: JR, TM, SM; Obtain and maintain biological material: TM, JR, DT; DNA preparation: JR, RL; Funding: JR, CE, SM; Data analysis: SM; Drafting MS: SM; Writing and revision: All.

## Supplementary Material

**Supplementary Table 1.**
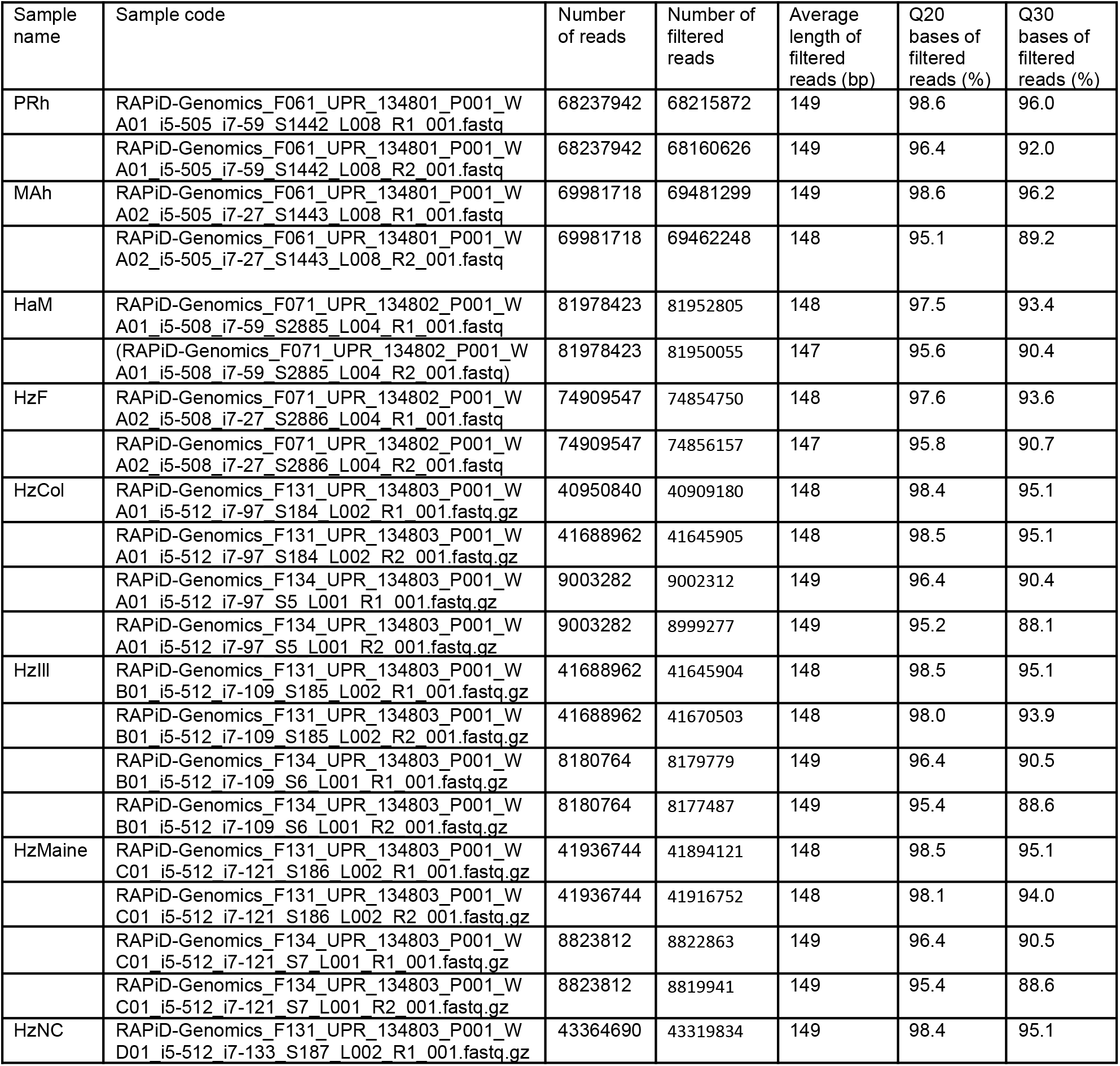
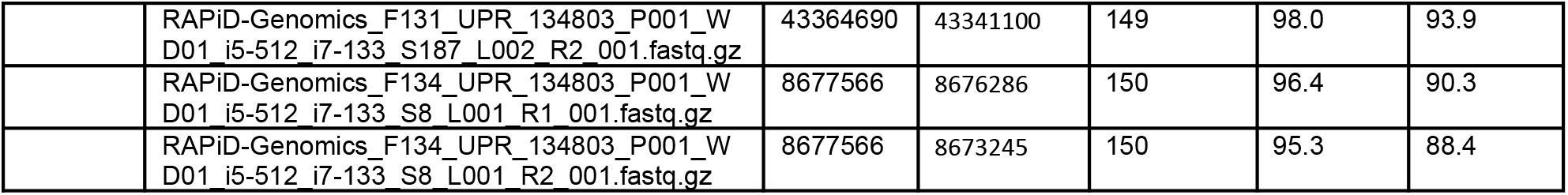
Sequencing statistics. Fastq files corresponding to this table can be found in the SRA under BioProject ID: PRJNA973566 or SRA submission number SUB13362524. The latter four genomes were sequenced in two separate lanes. After filtration and repair, reads from the two lanes were merged before mapping.

